# Enabling regiospecific di-halogenation in one-pot reactions using an engineered single-component flavin-dependent tryptophan halogenase

**DOI:** 10.1101/2025.11.10.687318

**Authors:** Hao Li, Jian-Wen Huang, Si Dai, Deyi Feng, Jiangli Liu, Nan Zhang, Yaojie Guo, Chun-Chi Chen, Rey-Ting Guo

**Affiliations:** School of Life Sciences, Hubei University, Wuhan, 430062, PR China; Zhejiang Key Laboratory of Medical Epigenetics, Hubei Hongshan Laboratory, Department of Immunology and Pathogen Biology, School of Basic Medical Sciences, Hangzhou Normal University, Hangzhou, 311121, PR China

**Keywords:** Halogenase, bromination, iodination, AetF, crystal structure, enzyme engineering, one-pot reaction

## Abstract

Installation of halogen atoms to organic compounds through flavin-dependent halogenases (FDHs) catalyzed reactions has emerged as an attractive strategy in organic biochemistry and synthetic biology. A cyanobacterial FDH termed AetF that contains FDH and flavin reductase in the same polypeptide chain could become a useful tool enzyme as the need of including a reductase is omitted. Notably, AetF exploits a unique substrate-interaction network to consecutively brominate tryptophan (Trp) on C5 and then C7 to generate di-brominated Trp. In this study, structure-based engineering is used to eliminate the C7 halogenation capacity of AetF. The resulting variant AetF-AIF that contains three residue substitutions mainly catalyzes bromination or iodination on C5 of Trp. We also show that combining AetF-AIF in tandem with the wild type AetF enables the generation of heterogeneously halogenated Trp in a one-pot reaction. The final products, 5-Br-7-I-Trp or 5-I-7-Br-Trp depending on the addition order of halides, account for a high ratio in the reaction mixture without additional purification step. These results display the potentials of single-component Trp-FDHs in a wide application of the halogenation reactions.

**Table of Content Graphic:** A single-component flavin-dependent halogenase that catalyzes di-bromination of tryptophan was engineered as a mono-halogenase and coupled in a one-pot reaction for the synthesis of heterogeneously halogenated tryptophan.

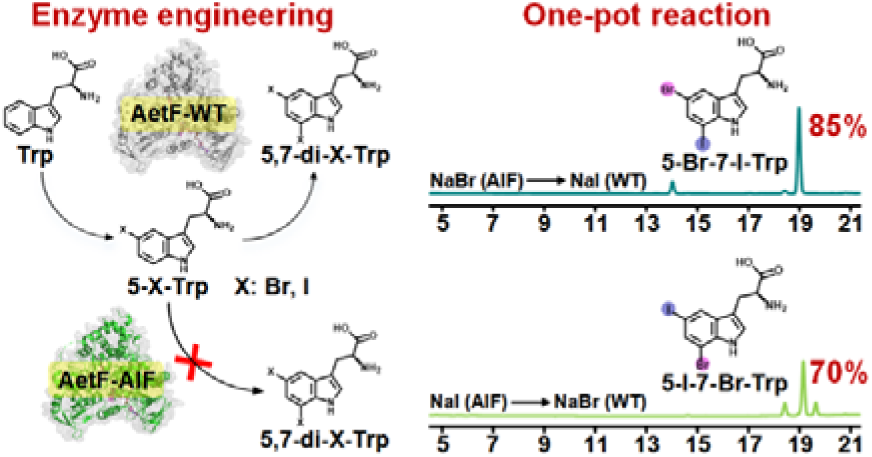

## Introduction

Organohalogens are present in many pharmaceutical and agrochemical products and are used in all sectors of the chemical industry as synthetic intermediates. Installation of halogen atoms could have profound effects in improving the bioactivity and physicochemical properties of small molecules. For instance, halogen introduction can be used to optimize lead drugs to increase the bioavailability and drug-target binding affinity without significantly compromising other interactions to the target [1,2]. In addition, halogen atoms may serve as chemical linchpins to facilitate additional derivatization of the molecular scaffold. Incorporation of halogens with chemical approaches typically requires toxic catalysts or metals and harsh condition, and the control of regiospecificity remains highly challenging. Therefore, enzyme-mediated halogenation that can operate a highly regiospecific manner under benign conditions without involving hazardous reagents has emerged as an attractive strategy [3].

A plethora of halogenated natural products and the halogenases in their biosynthetic routes have been identified [4,5]. Among these enzymes, flavin-dependent halogenases (FDHs), particularly those catalyze electrophilic halogenation toward substrates that are not anchored to a carrier, have attracted much attention [6]. The action of FDHs requires the reduced flavin and molecular oxygen to generate hydroperoxyl flavin intermediate, which oxidizes a halide ion to form a halogenating reagent hypohalous acid (HOX). The HOX is then guided to the substrate via a catalytic lysine that is located in the halfway of a tunnel that connects the FAD- and substrate-binding regions [7–9]. Extensive studies have been conducted with particular focus on flavin-dependent tryptophan halogenases (Trp-FDHs) that can install a halogen at either C5, C6 or C7 of the indole of Trp. Trp-FDHs that show usefulness in C-H functionalization, preparation of therapeutic peptide building blocks and modification of peptides are powerful synthetic biocatalysts [10–12]. It has also been demonstrated that Trp-FDHs-catalyzed halogenation enables selective chemoenzymatic C-H functionalization via methods such as Suzuki-Miyaura reaction [13,14]. However, further applications of canonical FDHs are hindered by low catalytic activity and instability [5,6,15]. Furthermore, the need to coupling a separate flavin reductase to supply the reduced flavin cofactor also complicates the establishment of the reaction system.

More recently, a class of “single-component” FDHs has been identified [16–19]. These enzymes fuse pyridine nucleotide-dependent flavin reductase and FDH on the same polypeptide chain [20–22], an advantageous feature in the application aspect relative to canonical FDHs owing to the omission of including a flavin reductase. The first documented single-component Trp-FDH is AetF from cyanobacterium *Aetokthonos hydrillicola*, which catalyzes consecutively bromination on C5 and then C7 of Trp to generate 5,7-di-Br-Trp [16,17]. The action of AetF indicates that both Trp and 5-Br-Trp can be used as a substrate. Recent structural reports demonstrate that the bi-specificity of AetF is conferred by the sophisticated designated substrate-binding pocket [21,22]. The indole ring of Trp and 5-Br-Trp lies in the same plane but flips around an axis formed by C3 and C6. The flipping allows the carbon-to-be-halogenated to be oriented toward the catalytic residue K258 and within the hydrogen bond distance to the putative general base E200, such that the halogenation site on two substrates is switched from C5 to C7 [21]. AetF possesses higher coupling rate and thus higher catalytic efficacy compared with canonical Trp-FDHs [18,23]. It has been suggested that the flavin oxygen adduct intermediates formed within AetF are stable in the absence of substrate, which minimizes unnecessary substrate consumption and the leakage of HOX [23].

Modulating the substrate selectivity and regioselectivity of canonical FDHs through engineering has been documented [24–28], and the capability of catalyzing a consecutive halogenation in a defined order makes AetF an attractive target to be engineered to become a regiospecific monohalogenase [22]. When conducting Ala substitution to investigate the role of the substrate-binding residues in our previous study, we found that substituting S523, a residue that binds the carboxyl group of the substrate, to Ala leads to 2.5-fold increase in 5-Br-Trp production and very low di-brominated product. In the present study, we focus on improving the performance of variant S523A of AetF through structure-based engineering and eventually obtained a triplet variant that shows strict C5-specific bromination activity. Notably, combining the variant in tandem with the wild type AetF enables the site-specific differentially di-halogenated Trp in a one-pot reaction.

## Materials and Methods

### Preparation of mutagenesis and recombinant proteins

The plasmid pET-32a/AetF that contains the AetF-encoding gene (GenBank no. WP_208344498.1) was prepared as described in our previous study [21]. The plasmids that encode variant AetF were constructed by polymerase chain reaction-based site-directed mutagenesis with plasmid pET-32a/AetF as a template. The mutagenic oligonucleotides used for the construction of variants were listed in **Table S1**. All plasmids were verified by direct sequencing.

The recombinant proteins of wild type and variant AetF were expressed and purified as previously described [21]. Briefly, the plasmids were transformed to *Escherichia coli* BL21(DE3), cultured in LB medium at 37 °C and induced by 0.3 mM isopropyl-β-□-thiogalactopyranoside when the optical density at 600 nm reached 0.6-0.8. Cells were collected after 16-20 h and centrifuged at 6,000 × g for 10 min and re-suspended in a lysis buffer containing 25 mM Tris-HCl (pH 7.5), 300 mM NaCl and 20 mM imidazole. After disruption with a French press, the debris was removed by centrifugation at 17,000 × g for 1 h and the resulting supernatant was loaded onto a nickel nitrilotriacetic acid agarose (Ni-NTA) affinity column. The target proteins were eluted using an imidazole gradient from 20 mM to 500 mM. Fractions containing AetF were collected, dialyzed against a buffer containing 25 mM Tris-HCl (pH 7.5) and 300 mM NaCl, and subjected to tobacco etch virus protease digestion to remove the thioredoxin and His_6_ tag. The untagged AetF proteins were obtained by passing through a Ni-NTA column again to remove the His-tagged portion. The protein purity (> 95 %) was verified by SDS-PAGE analysis and the concentration was determined by BCA assay.

### Crystallization and structure determination

The protein crystals were grown by using the previously reported crystallization conditions that 2 μL protein (10 mg mL^-1^) was mixed with 2 µL reservoir solution (25-30 % PEG 6000 and 100 mM Tris, pH 8.0) in 24-well Cryschem plates [21]. To obtain the complex structures, crystals were soaked with Trp for 0.5 h prior to the data collection. The X-ray diffraction datasets of AetF-S523A/Trp and AetF-AI/Trp were collected on beam line TPS-07A (λ = 0.97621 Å) and that of AetF-AIF/Trp on beam line TLS-15A (λ = 1.00000 Å) of the National Synchrotron Radiation Research Center (NSRRC, Hsinchu, Taiwan). All datasets were processed by using HKL2000 [29]. The complex structures were solved by the method of molecular replacement by using the complex structure of AetF/FAD (PDB ID, 8JZ2) as a search model. Subsequent model adjustment and refinement were conducted by using Refmac5 and Coot [30,31], with 5 % of randomly selected reflections set aside for calculating *R*_free_ as a monitor of model quality. All protein structure figures were prepared using the PyMOL program (http://pymol.sourceforge.net/). Data collection and refinement statistics are summarized in **Table 1**.

**Table 1.** Data collection and refinement statistics of crystal structures of variant AetF in complex with Trp.

|  | AetF-S523A/Trp | AetF-AI/Trp | AetF-AIF/Trp |
| --- | --- | --- | --- |
| PDB code | 9VWV | 9VWW | 9VWX |
| **Data collection** |  |  |  |
| Space group | $P2_12_12_1$ | $P2_12_12_1$ | $P2_12_12_1$ |
| Unit cell |  |  |  |
| a, b, c (Å) | 60.1, 75.0, 143.2 | 60.2, 75.4, 143.8 | 61.5, 74.9, 142.2 |
| $\alpha, \beta, \gamma$ (°) | 90, 90, 90 | 90, 90, 90 | 90, 90, 90 |
| Resolution (Å)<sup>a</sup> | 25.00-1.81<br>(1.87-1.81) | 25.00-2.32<br>(2.40-2.32) | 25.00-2.28<br>(2.36-2.28) |
| No. of observed reflections | 59211 (5595) | 28435 (2865) | 29977 (2959) |
| Redundancy | 4.2 (4.1) | 4.4 (4.3) | 5.2 (5.2) |
| Completeness (%) | 98.3 (94.1) | 98.1 (99.9) | 98.8 (99.1) |
| Average $I/\sigma$ (I) | 23.9 (2.2) | 18.2 (2.1) | 21.1 (3.0) |
| $R_{\text{merge}}$ (%)<sup>b</sup> | 5.4 (49.4) | 9.2 (61.7) | 8.7 (89.4) |
| CC 1/2 | 0.993 (0.905) | 0.996 (0.868) | 0.997 (0.865) |
| **Refinement<sup>c</sup>** |  |  |  |
| $R_{\text{work}}$ (%) | 19.0 (34.2) | 21.8 (30.8) | 19.4 (30.8) |
| $R_{\text{free}}$ (%) | 25.4 (32.8) | 29.5 (40.8) | 23.2 (28.5) |
| r.m.s.d. bonds (Å) | 0.008 | 0.006 | 0.007 |
| r.m.s.d. angles (°) | 1.536 | 1.440 | 1.510 |
| **Ramachandran statistics** |  |  |  |
| Most favored (%) | 96.5 | 95.3 | 96.3 |
| Allowed (%) | 3.5 | 4.7 | 3.7 |
| Outliers (%) | 0 | 0 | 0 |
| **Average B-factor (Å<sup>2</sup>)** |  |  |  |
| Protein/atoms | 46.1/5213 | 60.2/5202 | 54.8/5188 |
| Water/atoms | 46.3/312 | 51.2/83 | 45.8/155 |
| Ligand/atoms | 41.4/83 | 53.6/68 | 47.4/83 |

<sup>a</sup> Values in parentheses are for the highest resolution shell.
<sup>b</sup> $R_{\text{merge}} = \sum_{hkl} \sum_i |I_i(hkl) - \langle I(hkl) \rangle| / \sum_{hkl} \sum_i I_i(hkl)$ , in which the sum is over all the $i$ measured reflections
with equivalent miller indices $hkl$ ; $\langle I(hkl) \rangle$ is the averaged intensity of these $i$ reflections, and the
grand sum is over all measured reflections in the data set.
<sup>c</sup> All positive reflections were used in the refinement.

### Enzyme activity measurement

The typical reactions (200 μL) containing 0.65 μM AetF, 20 mM halide, 1 mM NADPH, 100 μM FAD, 0.5 mM Trp and 10 % (v/v) glycerol in 25 mM Tris-HCl buffer (pH 8.0) were incubated at 30 °C for 30 min. The reaction was terminated by adding 200 μL methanol and then analyzed by a high-performance liquid chromatography system (HPLC, Shimadzu LC-20AD, Japan) equipped with an SPD-M20A photodiode array detector. Separation was conducted by using the InerSustain C18 column (4.6 × 250 mM, 5 μm) with solvent A (double distilled water) and solvent B (methanol) with a linear gradient of 0-3 min 25 % B, 3-20 min 25-100 % B and 20-23 min 100 % B in a flow rate of 1.0 mL min^-1^. The relative activity of each variant was determined by the amounts of 5-Br-Trp and 5,7-di-Br-Trp, which were presented as a percentage of 5-Br-Trp produced by the wild type AetF as:

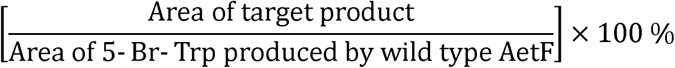

All assays were performed in triplicate and the results were presented as average ± SD.

For the detection of 7-Br-Trp, HPLC system coupled with a Obelisc R column (4.6 × 150 mM, 5 μm) eluted with solvent A (double distilled water) and solvent B (acetonitrile) in a flow rate of 1.0 mL min^-1^ were applied, employing gradients from 0-3 min 5 % B, 3-8 min 5-10 %, 8-15 min 10-30 % B, 15-25 min 30-95 % B and 25-35 min 5 % B to re-equilibrate the column.

### HOBr leakage assessment

The leaked HOBr was captured by using D-luciferin. 200 μL reactions containing 0.65 μM enzyme, 20 mM NaBr, 1 mM NADPH, 100 μM FAD, 0.5 mM D-luciferin in the absence or presence of 0.5 mM Trp and 10% (v/v) glycerol in 25 mM Tris-HCl buffer (pH 8.0) were incubated at 30 °C for 1 h. The reaction was terminated by adding 200 μL methanol and analyzed by HPLC system (Shimadzu LC-20AD, Japan) equipped with a SPD-M20A photodiode array detector at 330 nm. Separation was conducted by using the InerSustain C18 column (4.6 × 250 mM, 5 μm) with solvent A (double distilled water) and solvent B (methanol) with a linear gradient of 0-3 min 20% B, 3-15 min 20-50% B, 15-21 min 50-100% B and 21-23 min 100% B in a flow rate of 1.0 mL min^-1^. All assays were performed in triplicate and the results were presented as average ± SD.

### Kinetic assay

For the kinetic studies of wild type AetF and AetF-AIF when using Trp as a substrate, the reaction mixtures (200 μL) containing 0.26 μM enzyme, 20 mM NaBr, 1 mM NADPH, 100 μM FAD, 5 to 150 μM Trp and 10 % (v/v) glycerol in 25 mM Tris-HCl buffer (pH 8.0) were incubated at 30 °C. The samples were subjected to HPLC analysis following the abovementioned procedures. All assays were performed in triplicate and the results were presented as average ± SD. The enzyme reaction rate was calculated by the integral area of Trp:

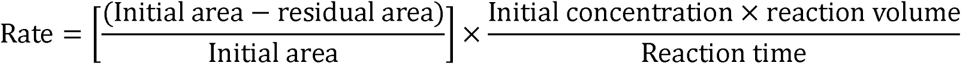

The kinetic parameters *K*_M_ and V_max_ were determined by fitting the reaction rate data to the Michaelis-Menten function of OriginPro 9.1.0 (OriginLab Corporation, Northampton, MA, USA).

### NADPH oxidation assay

The mixtures (200 μL) containing 1.3 μM enzyme, 20 mM NaBr, 0.2 mM NADPH and 10 % (v/v) glycerol in 25 mM Tris-HCl buffer (pH 8.0) were assembled. The reactions were initiated by adding varying concentrations of Trp (50, 100, 200 or 500 μM) in 96-well microplates. The NADPH oxidation was monitored through the changes of 340 nm by using SpectraMax M2 for 30 minutes at 30°C.

### One-pot reaction

One-pot synthesis of 5-Br-7-I-Trp (scheme A): (□) The reaction (200 μL) containing 1.3 μM AetF-AIF, 2 mM NaBr, 1 mM NADPH, 100 μM FAD, 0.5 mM Trp and 10 % (v/v) glycerol in 25 mM Tris-HCl buffer (pH 8.0) was incubated at 30 °C for 1 h. (□) The mixture was supplemented with 1.3 μM wild type AetF, 20 mM NaI, 0.5 mM NADPH and 50 μM FAD, and incubated at 30°C for another 1 h. One-pot synthesis of 5-I-7-Br-Trp (scheme B): (□) The reaction (200 μL) containing 3.2 μM AetF-AIF, 1 mM NaI, 1 mM NADPH, 100 μM FAD, 0.2 mM Trp and 10 % (v/v) glycerol in 25 mM Tris-HCl buffer (pH 8.0) was incubated at 30 °C for 1 h. (□) The mixture was supplemented with 3.2 μM wild type AetF, 50 mM NaBr, 0.5 mM NADPH and 50 μM FAD, and incubated at 30°C for another 1 h. The reaction mixtures were analyzed at 280 nm by using the HPLC analysis method described above. The percentage of each component in the mixture was calculated as:

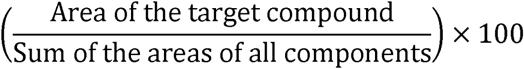

In addition, UHPLC-DAD-HRESIMS data of the products were collected in a negative ion mode by an Agilent Infinity II 1290 UHPLC-6546 QTOF MS system equipped with a diode array detector (DAD) as previously described [32].

## Results and discussion

### Crystal structure of AetF-S523A

Before conducting engineering, the crystal structure of AetF-S523A in complex with Trp was resolved (**Table 1**). The electron density maps corresponding to Trp were clearly observed in the substrate-binding pocket (**Fig. 1A**). The binding pose and the interaction network of Trp to the enzyme are identical to those of the wild type enzyme (**Fig. 1B**). This indicates that removing S523-mediated polar interaction poses minimal influence to the binding of Trp. Noteworthily, we failed to obtain the complex structure of AetF-S523A and 5-Br-Trp despite of great efforts, which should be attributed to the low affinity of the variant enzyme to the compound. This also indicates that the S523-mediated interaction is crucial to the binding of 5-Br-Trp, such that the Ala substitute is incapable of accepting the substrate that shows a flipped pose relative to Trp and displays low dibrominated Trp production activity as previously proposed [22]. It should be noted that AetF-S523A still produces trace amounts of 5,7-di-Br-Trp (**Fig. 1D**).

**Figure 1.**
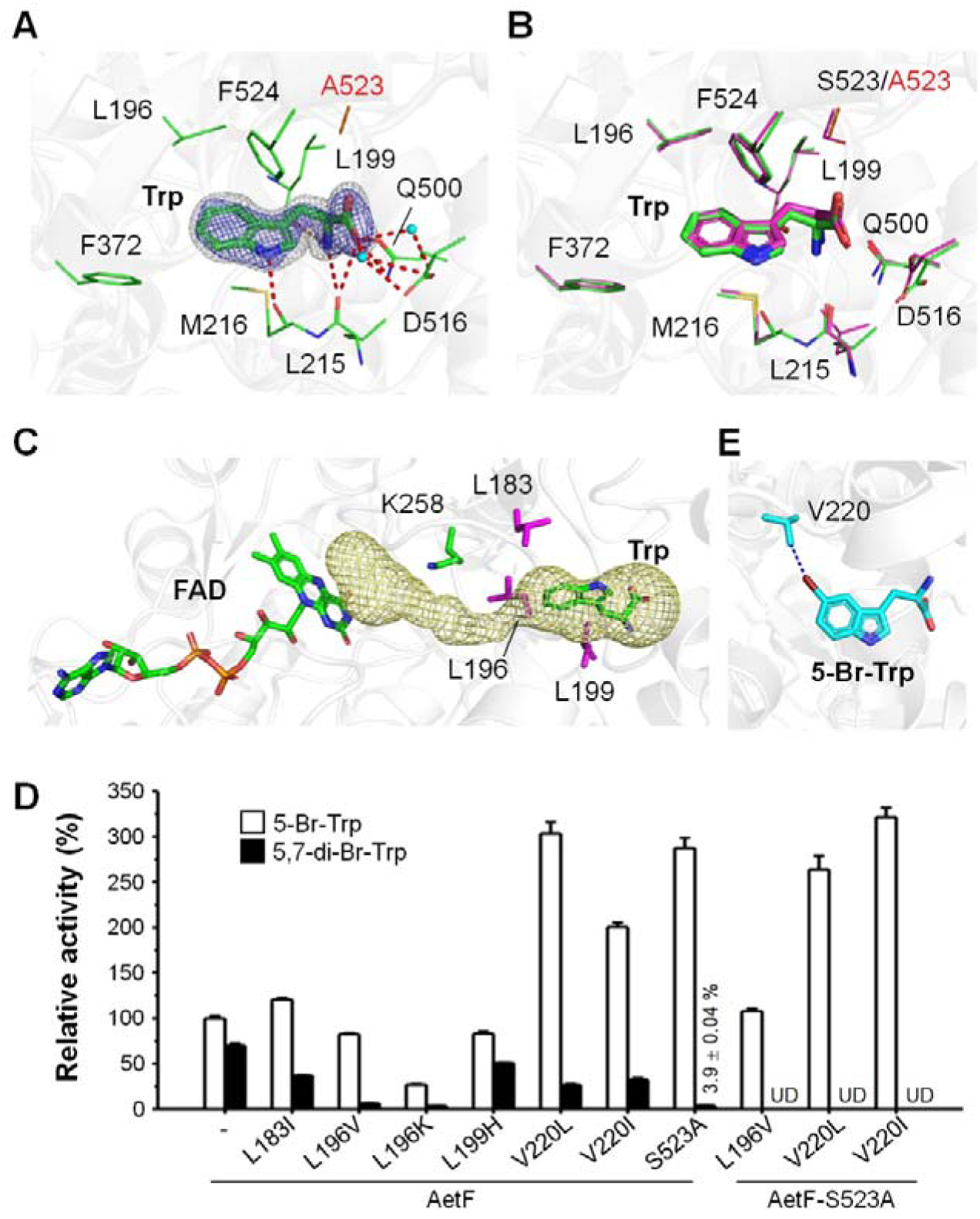
Engineering AetF-S523A. (**A**) The residues that comprise the Trp-interaction network in the crystal structure of AetF-S523A/Trp (PDB ID, 9VWV) are shown in lines, with A523 colored in red. The 2*F*_o_-*F*_c_ electron density map (white mesh) and *F*_o_-*F*_c_ omit map (blue mesh) of Trp are contoured at 1.0 and 3.0 σ, respectively. Red dashed lines, distance < 3.5 Å. Cyan spheres, water molecules. (**B**) Structural superimposition of AetF-S523A/Trp (green) and AetF/Trp (magenta; PDB ID, 8JZ3). The residues that constitute the Trp-interaction networks are shown as described in (**A**). (**C**) The putative tunnel transversing from FAD to Trp in AetF-S523A/Trp is displayed with pale-yellow bubble. The catalytic K258 (green) and several residues that surround the tunnel (magenta) are indicated. (**D**) The activity of variant AetF (0.65 μM) was determined by measuring the yield of 5-Br-Trp and 5,7-di-Br-Trp in 30 min, which are presented as a percentage of 5-Br-Trp produced by the wild type AetF. Triplicate assay was conducted and each sample were analyzed for three independent times. The results are presented as average ± SD. UD, undetectable. (**E**) The location of residue V220 that is adjacent to the brominated site of 5-Br-Trp in the crystal structure of AetF/5-Br-Trp (PDB ID, 8JZ4). Blue dashed line, 3.4 Å.

### Improving the performance of AetF-S523A

First, we aimed to modify the HOX transfer tunnel, a strategy that has been demonstrated to enhance the enzyme stability through lowering the leakage of HOX [33]. Prakinee et al. conducted mechanism-guided semi-rational approach and identified a variant that harbors a single substitution of a residue located between the catalytic Lys and the substrate. In analogy to the previous study, several residues that are located between K258 and the Trp-binding site were selected to construct a series of variants (**Fig. 1C**). As shown in **Fig. 1D**, variant L183I shows a moderate increase in the production of 5-Br-Trp and lower di-brominated product. The production of 5-Br-Trp of L196V was approximately 80 % compared with the wild type AetF, but much lower di-brominated product was yielded. The catalytic activity of two variants of L199 are decreased compared with the wild type enzyme. Variant L199H exhibits higher activity than L199K, but only retains 70 to 80 % activity compared with the wild type AetF. The attempts to modify the HOX tunnel did not improve the performance of AetF, which might owe to the fact that AetF inherently exhibits very low HOX leakage compared with canonical FDHs [15]. Consistent with the previous report, the HOBr leakage rate assessed by using D-luciferin as an indicator is low in AetF-catalyzed reaction (**Fig. S1**).

We also inspected the structure of AetF in complex with 5-Br-Trp and found that V220 is in close vicinity to the Br on C5 (**Fig. 1E**). We thus substituted V220 with bulkier residues including Ile and Leu in an attempt to reduce the enzyme’s capability in binding 5-Br-Trp. Both variant of V220 exhibit higher production of 5-Br-Trp and lower production of 5,7-di-Br-Trp (**Fig. 1D**). We thus introduced mutations including L196V, V220I and V220L to AetF-S523A. Unlike AetF-S523A, no di-brominated Trp was detectable in all doublets with AetF-V220I/S523A, hereinafter termed AetF-AI, being the most effective variant, whose 5-Br-Trp production is 3.2-fold higher than the wild type enzyme (**Fig. 1D**).

### The substrate specificity of AetF-AI

Given that the goal is transforming AetF to a C5 mono-halogenase, the regiospecificity of AetF-AI was examined cautiously. We assembled reactions that contains 5-Br-Trp as a substrate and investigated the reaction products in prolonged incubation time. As shown in **Fig. 2A**, more than 50 % of 5-Br-Trp was converted to di-brominated Trp by the wild type AetF within 30 min and all substrate was consumed in two hrs. Compared with the wild type enzyme, AetF-AI exhibits much lower activity against 5-Br-Trp but approximately 25 % substrate conversion occurred in 8 hrs, suggesting that the variant still possesses C7 halogenase activity.

**Figure 2.**
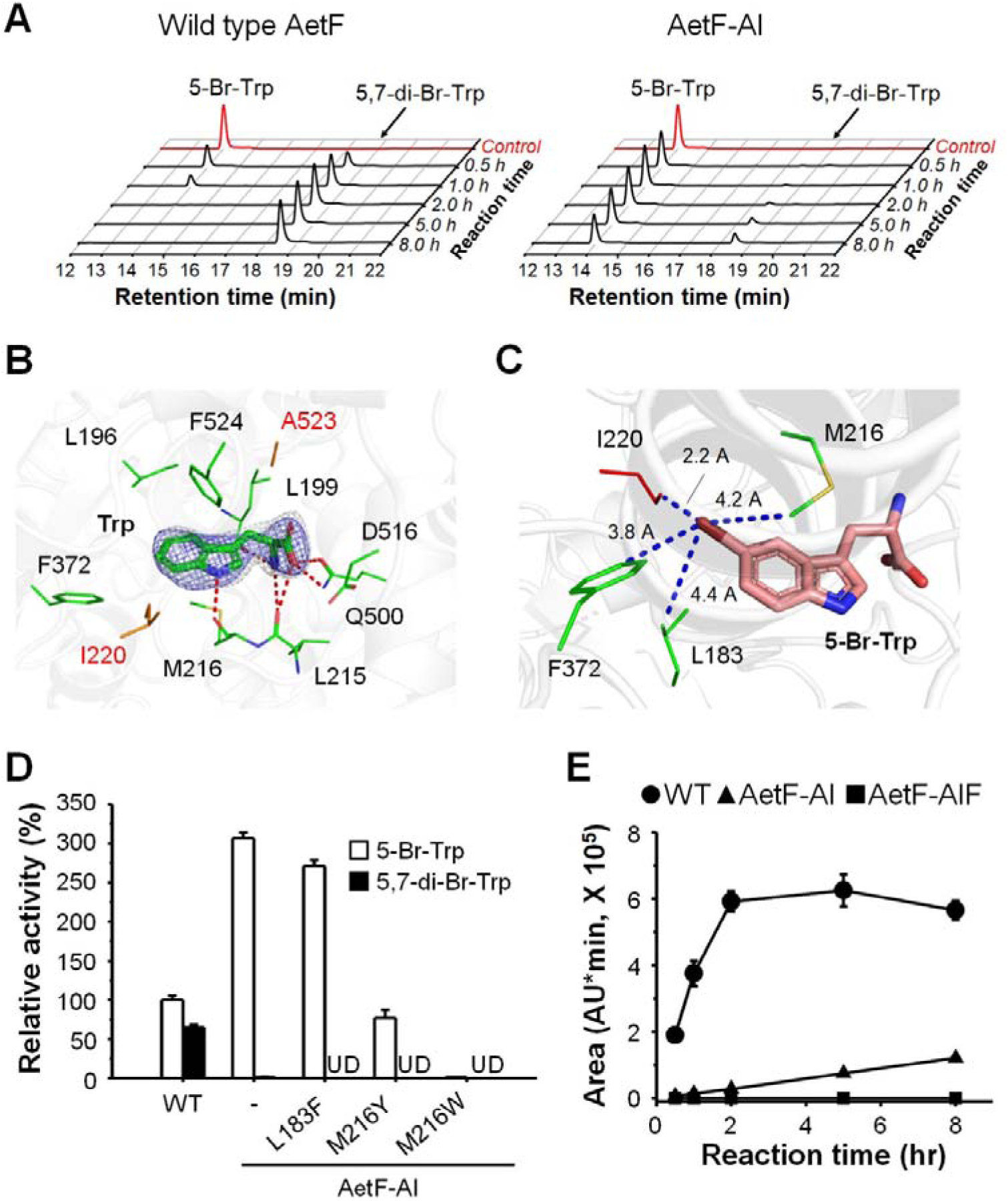
Reducing di-bromination activity of AetF-AI. (**A**) The representative HPLC chromatograms of reaction products of 5-Br-Trp catalyzed by wild type AetF and AetF-AI. Control, reaction without enzyme. (**B**) The Trp-interaction residues in the crystal structure of AetF-AI/Trp (PDB ID, 9VWW) are shown in lines, with A523 and I220 colored in red. The electron density maps of Trp are contoured and displayed as described in Fig. 1A. Red dashed lines, distance < 3.5 Å. (**C**) Model of 5-Br-Trp-binding mode in AetF-AI. The 5-Br-Trp (salmon stick) from the structure of AetF/5-Br-Trp (PDB ID, 8JZ4) is modeled to the structure of AetF-AI/Trp. Several Br-surrounding residues are shown in lines. Dashed lines measure the shortest distances between the residues and the Br with the length indicated. (**D**) The activity of variant AetF was determined and presented as described in Fig. 1D. All assays were performed in triplicate and the results were presented as average ± SD. UD, undetectable. (**E**) The 5,7-di-Br-Trp production of wild type and variant AetF at indicated time points. Triplicate assay was performed and the results were presented as average ± SD. The representative HPLC chromatograms of each group are displayed in **Fig. S2**.

We thus continued to modify AetF-AI in an attempt to eliminate the C7 halogenation activity. The crystal structure of AetF-AI in complex with Trp was solved to reveal the substrate-binding mode to facilitate the subsequent rationale design (**Table 1 and Fig. 2B**). The substrate was found to bind to the same location as those in the WT and AetF-S523A, suggesting that the substitution of V220 with Ile marginally, if any, influences the substrate-binding pose (**Fig. S2**). As the case of AetF-S523A, we failed to obtain the complex structure of AetF-AI and 5-Br-Trp. Therefore, we assessed the putative binding pose of 5-Br-Trp by modeling 5-Br-Trp from the AetF/5-Br-Trp complex structure to the crystal structure of AetF-AI/Trp (**Fig. 2C**). From this model, the side chain of I220 is distant from the Br on 5-Br-Trp by only 2.2 Å, a distance that may exclude the binding of 5-Br-Trp.

Based on these analyses, we considered that additional steric hindrance in the vicinity of Br on the 5-Br-Trp may assist to reduce the binding of 5-Br-Trp so as the production of 5,7-di-Br-Trp of AetF-AI. We thus focused on modifying M216, F372 and L183 and eventually obtained recombinant proteins of triple variants including AI-L183F, AI-M216Y and AI-M216W and had their catalytic activity tested. Variants containing M216 substitutions exhibit much lower activity than AetF-AI and the wild type enzyme thus were not further investigated (**Fig. 2D**). AI-L183F, termed AetF-AIF hereinafter, shows higher production of 5-Br-Trp compared with wild type AetF (271 ± 7.7 %). Despite the 5-Br-Trp production level of AetF-AIF is slightly lower than AetF-AI (271 ± 7.7 % vs. 306.7 ± 6.9 %) (**Fig. 2D**), this variant produces no detectable di-brominated product from 5-Br-Trp in reaction incubated as long as 8 hrs (**Fig. 2E and Fig. S2**). To further validate the regiospecificity AetF-AIF, we also investigated whether wild-type and variant AetF could produce 7-Br-Trp and found that no 7-Br-Trp can be detected (**Fig. S4**). Therefore, AetF-AIF was tentatively defined as a C5 Trp-FDH, which was subjected to the subsequent investigations.

### Kinetic and structural investigations of AetF-AIF

Next, kinetic experiments were conducted to assess the substrate-binding affinity and catalytic efficacy of AetF-AIF. As shown in **Table 2**, AetF-AIF exhibits higher *K*_M_ and *k*_cat_ against Trp compared with the wild type enzyme, suggesting that the triple mutations render AetF higher catalytic rate and lower Trp-binding capacity. A recent study demonstrates that the presence of Trp could accelerate the rate of NADPH oxidation of AetF [23]. We thus also examined the oxidation NADPH rate of wild type AetF and AetF-AIF when various concentration of Trp was supplemented in the reactions. As shown in **Fig. 3A**, the NADPH oxidation rate reaches plateau at a concentration of 100 to 200 μM Trp in the reaction containing the wild type AetF, whereas at least 200 μM Trp was required for AetF-AIF to approach a similar level. These results suggest that higher concentration of Trp is required to reach the maximal catalytic efficiency in AetF-AIF than in wild type AetF, which also substantiates the results of kinetic study. Structural investigations indicate that Trp adopts the same binding pose in wild type enzyme and AetF-AIF (**Fig. 3B**). Intriguingly, we found that D-luciferin conversion rate in the absence of Trp is reduced in AetF variants compared with the wild type enzyme (**Fig. S1**), which may be attributed to the narrowed substrate-binding pocket that hinders the entry of D-luciferin.

**Figure 3.**
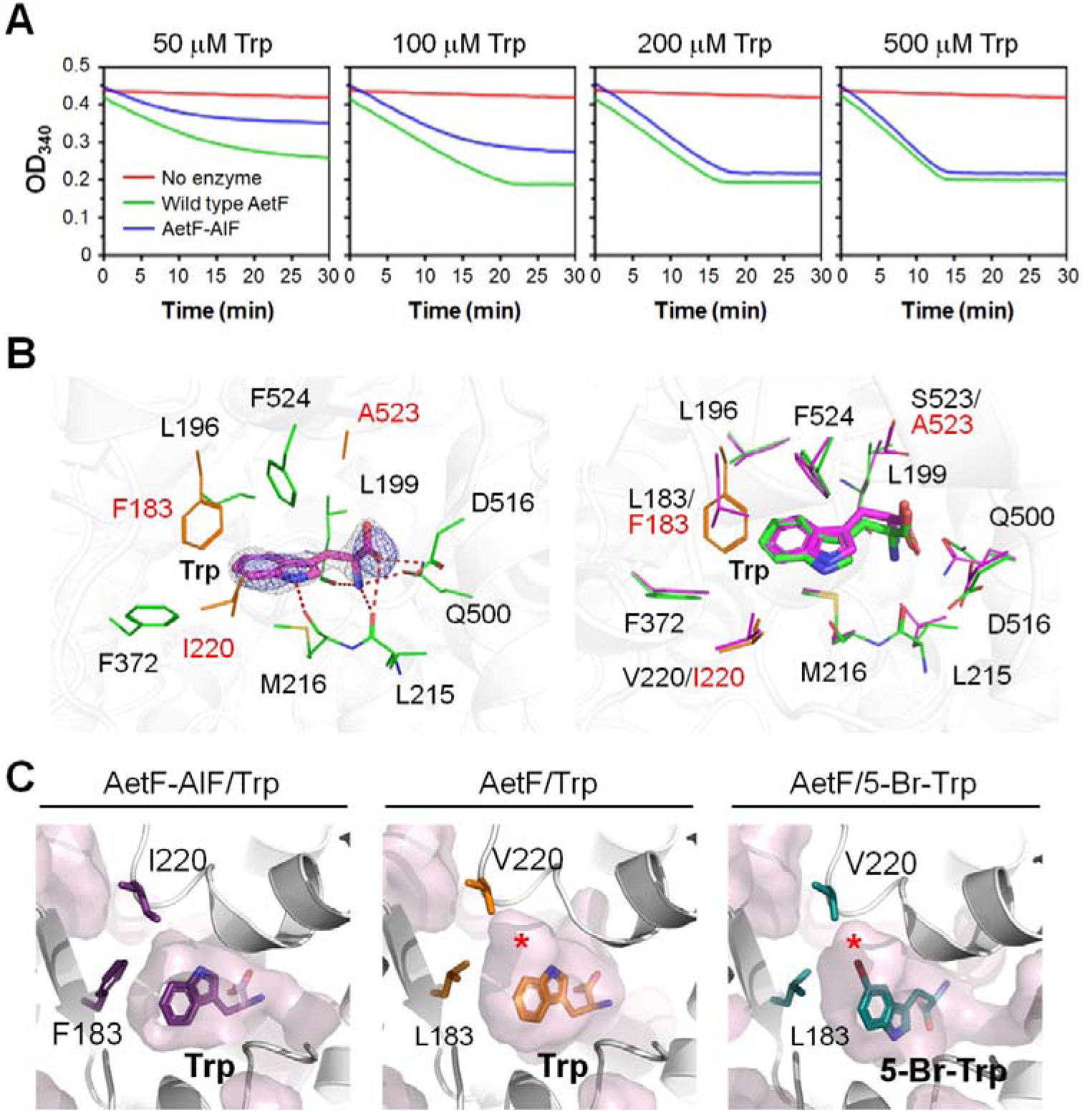
NADPH oxidation rate and substrate-binding mode of AetF-AIF. (**A**) The NADPH oxidation rates of wild type AetF and AetF-AIF when using Trp as a substrate. (**B**) Left: the residues that comprise the Trp-binding site in the crystal structure of AetF-AIF/Trp (PDB ID, 9VWX) are shown in lines, with A523, I220 and F183 colored in red. The electron density maps of Trp are contoured and displayed as described in Fig. 1A. Red dashed lines, distance < 3.5 Å. Right: the superimposition of AetF-AIF/Trp (green) and AetF/Trp (magenta; PDB ID, 8JZ3). The residues that constitute the Trp-interaction networks are shown in lines. (**C**) The Trp-binding pocket in AetF-AIF/Trp and AetF/Trp (PDB ID, 8JZ3) and 5-Br-Trp-binding pocket in AetF/5-Br-Trp (PDB ID, 8JZ4) are shown in pink bubble models. Residues 220 and 183 are displayed in sticks and the bulge that houses the Br of 5-Br-Trp is indicated with asterisks.

**Table 2.** Kinetic study of wild type AetF and AetF-AIF*.

| | $V_{\max}$ ($\mu\text{M min}^{-1}$) | $K_M$ ($\mu\text{M}$) | $k_{\text{cat}}$ ($\text{min}^{-1}$) |
| --- | --- | --- | --- |
| Wild type AetF | $1.57 \pm 0.1$ | $21.17 \pm 1.72$ | $6.05 \pm 0.37$ |
| AetF-AIF | $2.52 \pm 0.1$ | $63.7 \pm 6.51$ | $9.7 \pm 0.39$ |

\*Trp is used as a substrate.

We solved the complex structure of AetF-AIF and Trp and found that the bulge that accommodates the Br on the 5-Br-Trp is missing AetF-AIF (**Table 1 and Fig. 3C**). Altogether, we conclude that blocking the space near C5 in 5-Br-Trp in the AetF/5-Br-Trp in addition to preventing the flipping of the substrate is required to achieve complete elimination of C7-halogenation activity of AetF-AIF.

### One-pot reactions to generate heterogeneously di-halogenated Trp

The emergence of AetF-AIF prompted us to examine the possibility of synthesizing Trp that carries different halogens in a one-pot reaction system. First, the capacity to chlorinate and iodinate Trp of AetF-AIF was examined. Consistent with the previously report, wild type AetF, as well as AetF-AIF, showed very low chlorination activity [9], such that trace amounts of mono-chlorinated Trp, suspected to be 5-Cl-Trp, were detected even when the reactions contain as high as 500 mM NaCl and were incubated for 5 hrs (**Fig. S5**). Compared with chlorination, AetF catalyzes iodination more effectively (**Fig. S6**). Similar to the bromination capacity, AetF-AIF catalyzes iodination with a strong regioselectivity on C5, yet trace amount of di-iodinated Trp was yielded (**Fig. S6**). These results indicate that AetF is prone to iodinate the C7 of Trp and/or 5-I-Trp, though the underlying mechanism that accounts for the preference of using various halogens in the reaction remains unclear.

Given the limited chlorination activity of AetF and AetF-AIF, we focused on conducting bromination and iodination to synthesize di-halogenated Trp in the following trials. AetF-AIF that catalyzes C5 halogenation is supplemented first, and then the wild type AetF is added as a C7 Trp-FDH to halogenate the C7 (**Fig. 4A**). Two schemes were designated depending on the order of adding NaBr or NaI, which were expected to afford 5-Br-7-I-Trp (scheme A) or 5-I-7-Br-Trp (scheme B). The concentration of the primary halogen was kept as low as possible to prevent unwanted halogenation catalyzed by the wild type AetF in the second step. On the other hand, higher concentration of the secondary halogen was used to ensure the complete halogenation in the second step.

**Figure 4.**
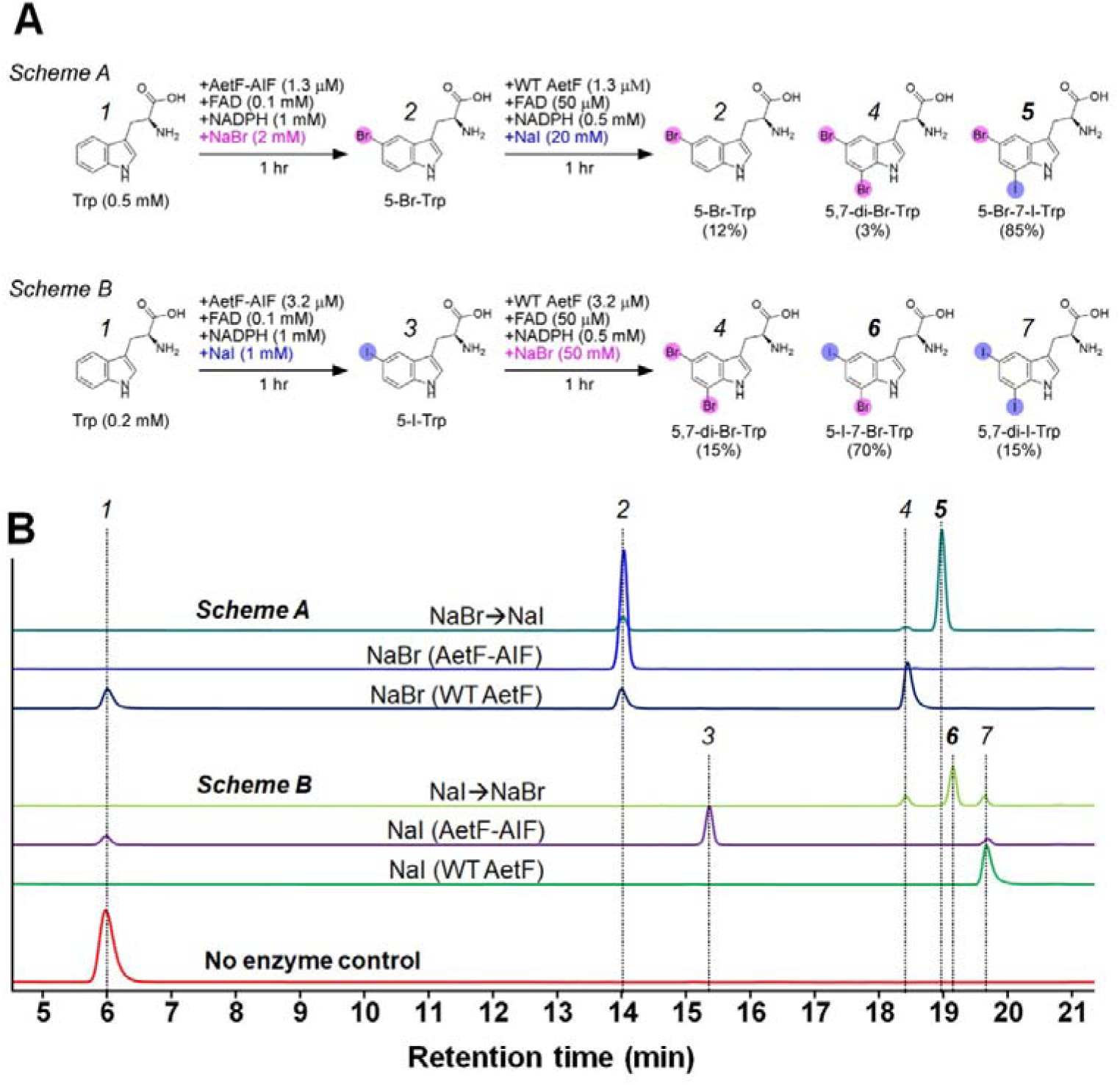
Synthesis of heterogeneously di-halogenated Trp in one-pot reactions. (**A**) The designation and the composition of the final product of two schemes. (**B**) The representative HPLC chromatograms of reaction mixtures yielded from two schemes described in (**A**). 1, Trp; 2, 5-Br-Trp; 3, 5-I-Trp; 4, 5,7-di-Br-Trp; 5, 5-Br-7-I-Trp; 6, 5-I-7-Br-Trp; 7, 5,7-di-I-Trp. The MS spectra of reaction mixtures from two schemes are shown in **Fig. S7** and **Fig. S8**.

In scheme A, Trp was converted to 5-Br-Trp by AetF-AIF in the first step and then iodinated on C7 by the wild type AetF to afford 5-Br-7-I-Trp (**Fig. 4B**). Three components including 5-Br-Trp, 5,7-di-Br-Trp and 5-Br-7-I-Trp were detected in the reaction mixture, with the target compound 5-Br-7-I-Trp accounting for 85 % and approximately 12 % of 5-Br-Trp that remains unreacted. Trace amount of 5,7-di-Br-Trp (3 %) was detected, which should be generated from 5-Br-Trp and NaBr retained in the reaction system by the wild type AetF that was supplemented in the secondary step. Intuitively, the undesirable bromination occurred in the secondary step could be prevented by reducing the concentration of NaBr supplemented in the reaction from the very beginning. However, lowering the concentration of NaBr resulted in incomplete transformation of Trp in the first step, which leads to additional products including 5-I-Trp and 5,7-di-I-Trp in the secondary step (data not shown). In scheme B, around 88% of Trp was converted to 5-I-Trp (85 %) and 5,7-di-I-Trp (15 %) by AetF-AIF in the first half of the reaction (**Fig. 4B**). As shown in **Fig. S6**, the transformation of Trp cannot be further elevated even in an elongated reaction time. Reducing the concentration of Trp or NaI also did not yield better results (data not shown). After adding the wild type AetF, 5-I-7-Br-Trp was generated as the main product, which accounts for 70 % of all components detected in the mixture. Notably, Trp retained from the first step served as a substrate of the wild type enzyme and was converted to di-halogenated Trp. Therefore, the end product of scheme B also contains 15 % of di-iodinated Trp and 15 % of di-brominated Trp.

The same results were observed when the volume of the reactions was magnified to 2 mL (**Fig. S9**), indicating that a tenfold amplification of the reaction system is viable. However, further scaling-up would require large amounts of cofactors and purified protein, which is not economically appealing for industrial applications. In view of this, other systems such as cell-based platform [34,35] or cross-linked enzyme aggregation method [36,37] should be opted for large-scale preparations.

Compared with AetF-AIF, utilization of AetF-S523A in the first half of the one-pot reaction increased the ratio of di-brominated Trp and lowered the ratio of 5-Br-7-I-Trp (from 85 % to 73 %) in the mixture of scheme A (**Fig. S10**). Utilization of AetF-S523A obviously impacted the reaction operated with scheme B, such that only 8 % of 5-I-7-Br-Trp was eventually yielded (70% in AetF-AIF-containing reaction) (**Fig. S10**). These results clearly indicate that the residual di-halogenated activity of AetF-S523, even low as around 3.9 % compared to the wild type enzyme as shown in **Fig. 1D**, could complicate the one-pot reaction outcome.

## Conclusion

In this study, we exploited rational design strategy to engineer a self-sufficient di-halogenase so that the variant enzyme catalyzes bromination or iodination primarily on C5 of Trp. Changing the regioselectivity of canonical FDHs have been documented [24–28], but the results often show compromised activity and mixed catalytic products. AetF-S523A, a variant that was identified in our previous study, exhibits around 3.9 % activity to produce di-brominate Trp compared with the wild type enzyme. However, even such level of C7-halogenation activity can lead to unwanted outcome in the one-pot reaction (**Fig. S10**). Therefore, elimination of the C7 halogenation activity of AetF should be necessary to grant this enzyme merits in further applications. In comparison, variant AetF-AIF shows highly specific regioselectivity and comparable activity to the wild type enzyme. We also show that combining AetF-AIF and the wild type AetF can achieve the synthesis of heterogeneously di-halogenated Trp in a one-pot reaction. The identity of each component in the reaction mixtures can be unambiguously determined through HPLC and mass spectrometry. To our knowledge, AetF-AIF is the first single-component Trp-FDH that exerts mono-bromination activity. In addition, the demonstrated one-pot reaction is the first reported system that could generate high purify of mixed-type halogenated compounds without additional purification step during the catalytic process [35,38]. Despite further improvements in the regioselectivity and catalytic efficacy of these Trp-FDHs should be required to optimize the reactions, it is considered that the presented results warrant the application potentials of AetF and the variants in synthetic biology. During the submission of our manuscript, Montua et al. published their work in using regio-complementary canonical Trp-FDHs and AetF to synthesize di- and tri-halogenated Trps using a combined cross-linked enzyme aggregation method [36,37]. Although the purity and the yield of the target compounds are lower than the present report and removing the catalytic entities in the preceding step is required, this system is attractive in terms of preparative scale-up reactions. It would be interesting to explore the efficacy of combining AetF and AetF-AIF by using such a platform.

## Supporting information

Mutagenesis oligonucleotides; D-luciferin transformation rate of wild type and variant AetF; structure superimpositions of AetF-AIF/Trp to AetF/Trp and AetF-AI/Trp; representative HPLC chromatograms of samples shown in **Fig. 2E**, chlorination and iodination reaction; extracted ion chromatograms and mass spectra of compounds in chlorination, iodination and one-pot reactions; scaled-up one-pot reactions.

## Acknowledgement

This work was supported by the National Key Research and Development Program of China (2021YFC2104000), Hubei Hongshan Laboratory (2022hszd030), the National Natural Science Foundation of China (32371307, 32271318 and 82341210); and the Interdisciplinary Research Project of Hangzhou Normal University (2024JCXK02). We thank NSRRC (National Synchrotron Radiation Research Center, Taiwan) for access to beam lines TPS-05A and TPS-07A that contributed for the synchrotron data collection.

