## Supplementary Information for "Enabling regiospecific di-halogenation in one-pot reactions using an engineered single-component flavin-dependent tryptophan halogenase"

**Affiliations**

**Supplementary Table**

**Table S1.** Mutagenesis oligonucleotides

| **Mutant** | **Sequence (5’→3’)** |
| --- | --- |
| L183I_F | TTTACAGCAATTGATCAGGAAGTGCAGGTT |
| L183I_R | TTCCTGATCAATTGCTGTAAAGCCATTGGT |
| L183F_F | TTTACAGCATTCGATCAGGAAGTGCAGGTT |
| L183F_R | TTCCTGATCGAATGCTGTAAAGCCATTGGT |
| L196V_F | CCGTTTACAGTGGATCAGCTGGAAAGCCCG |
| L196V_R | CAGCTGATCCACTGTAAACGGTTTGCCCAG |
| L199K_F | CTGGATCAGAAGGAAAGCCCGAATTTTCGT |
| L199K_R | CGGGCTTTCCTTCTGATCCAGTGTAAACGG |
| L199H_F | CTGGATCAGCACGAAAGCCCGAATTTTCGT |
| L199H_R | CGGGCTTTCGTGCTGATCCAGTGTAAACGG |
| M216Y_F | GATCGTCTGTACATGAGCCCGATTTATCCG |
| M216Y_R | CGGGCTCATGTACAGACGATCATACAGTTC |
| M216W_F | GATCGTCTGTGGATGAGCCCGATTTATCCG |
| M216W_R | CGGGCTCATCCACAGACGATCATACAGTTC |
| V220I_F | ATGAGCCCGATTTATCCGCGTACCGTTAAT |
| V220I_R | ACGCGGATAAATCGGGCTCATCATCAGACG |
| V220L_F | ATGAGCCCGCTTTATCCGCGTACCGTTAAT |
| V220L_R | ACGCGGATAAAGCGGGCTCATCATCAGACG |
| S523A_F | CGTCGCTTTGCCTTTAATGATATGCGTCAT |
| S523A_R | ATCATTAAAGGCAAAGCGACGGGCAGAATG |

^*^ The underlined nucleotides are mutation sites.

**Supplementary Figure**

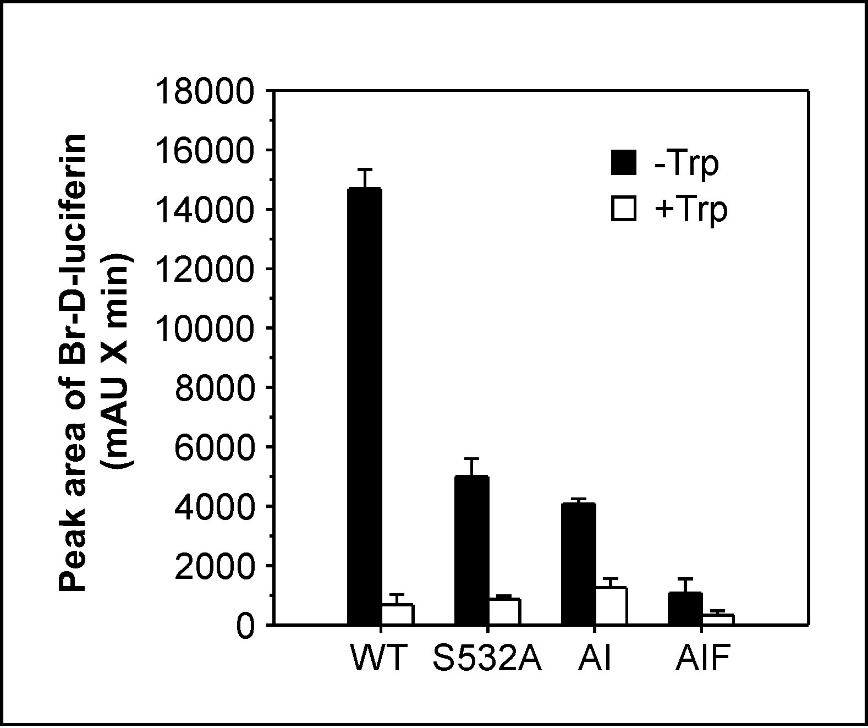

**Figure S1. Detection of HOBr leakage of AetF-catalyzed bromination reactions.** 200 μL reactions containing 0.65 μM wild type or variant AetF, 0.1 mM FAD, 1 mM NADPH, 20 mM NaBr, 0.5 mM D-luciferin in 20 mM Tris-HCl (pH 8.0) supplemented with 10 % (v/v) glycerol are incubated with or without 0.5 mM Trp at 30 °C for 1 h. The peak areas of Br-D-luciferin of each sample were recorded. Triplicate assays were performed and the results were presented as average ± SD. D-luciferin can be brominated by AetF (filled bars) and the reaction is profoundly suppressed in the presence of the natural substrate Trp (empty bars). For the wild type enzyme, the conversions of D-luciferin were calculated to approximately 0.3 % and 0.01 % in the absence and presence of Trp, respectively, comparable to the data reported by Phintha et al. {Phintha, 2025 #108}. Notably, the binding of D-luciferin in variant AetF is reduced compared with the wild type enzyme, a phenomenon supposed to be resulted from the altered substrate-binding pocket. As such, AetF-AIF that harbors a smallest substrate-binding pocket among all tested enzymes exhibits the lowest D-luciferin bromination capacity.

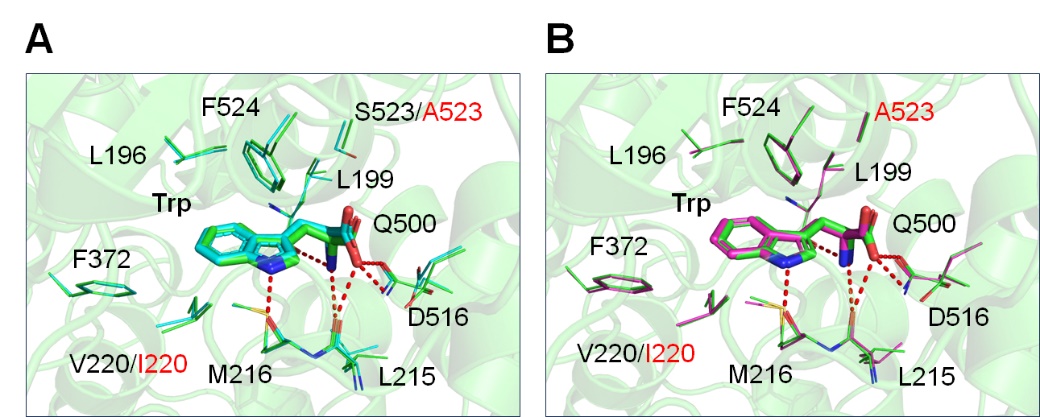

**Figure S2. Trp-binding modes in wild type and variant AetF.** Structural superimposition of AetF-AI/Trp (green; PDB ID, 9VWW) and (**A**) AetF/Trp (PDB ID, 8JZ3) or (**B**) AetF-AI/Trp (magenta; PDB ID, 9VWV). Residues that comprise the Trp-interaction networks are shown in lines, with the substituted ones indicated by red fonts. The overall structure of AetF-AI/Trp is shown in a cartoon model in two drawings and the Trp molecules in three structures in sticks. Red dashed lines, distance < 3.5 Å.

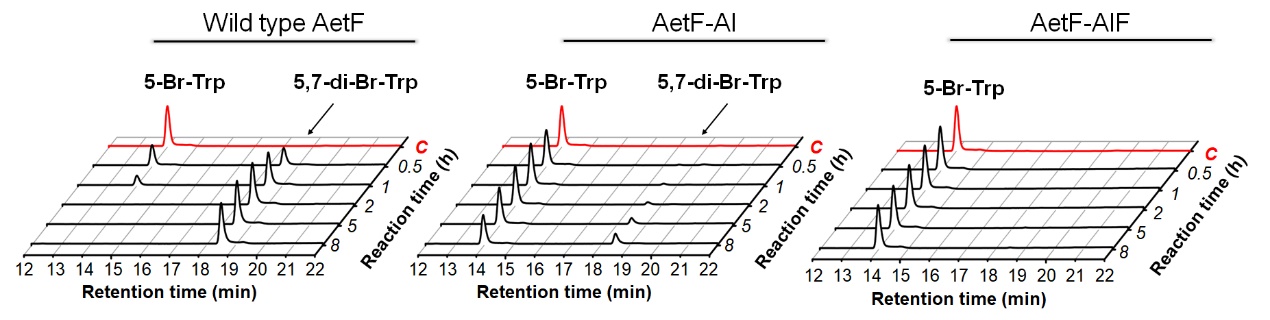

**Figure S3. Representative HPLC chromatograms of reaction products of wild type and variant AetF described in Fig. 2E.** C (control), reaction without enzyme.

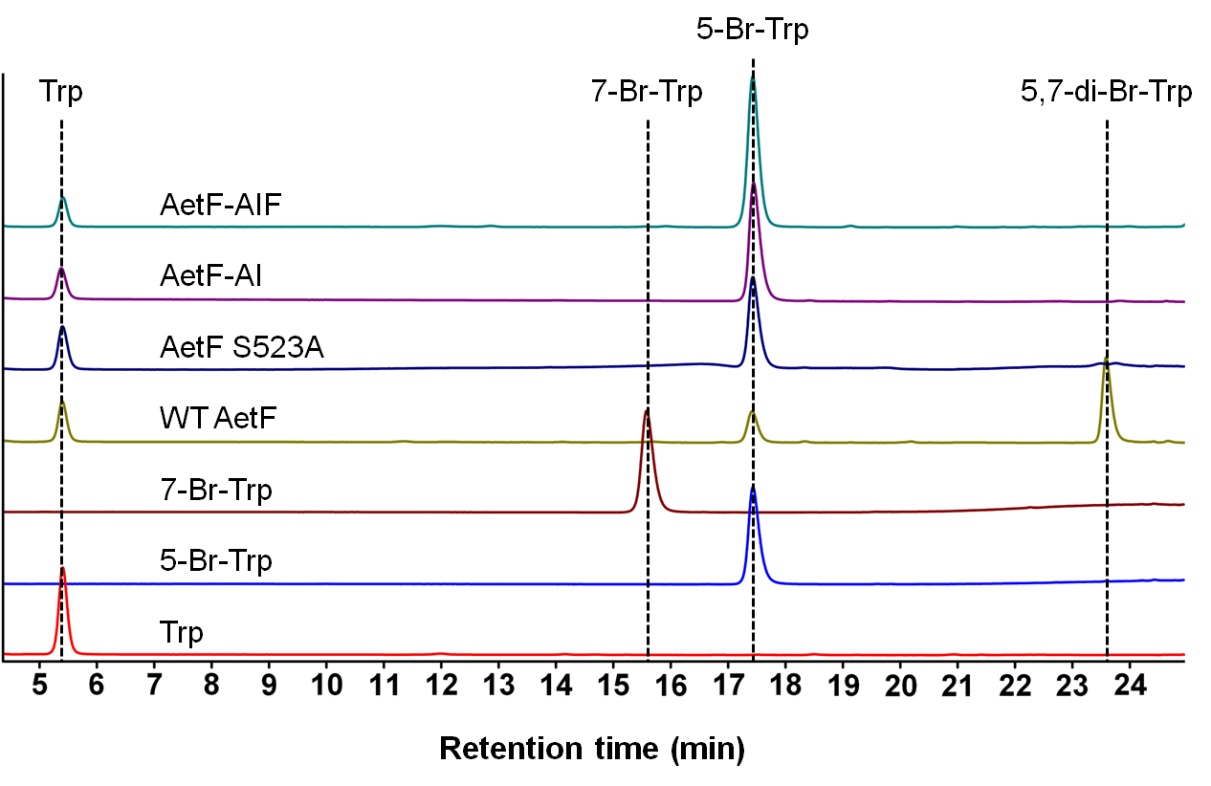

**Figure S4. The HPLC analyses of 7-Br-Trp in AetF-mediated reactions.** 200 μL reaction containing 1.3 μM AetF (wid type or variant), 20 mM NaBr, 1 mM NADPH, 100 μM FAD, 0.5 mM Trp and 10 % glycerol in 25 mM Tris-HCl buffer incubated at 30°C for 30 min and analyzed by HPLC coupled with Obelisc R column as described in Materials and Methods. The chromatograms of Trp, 5-Br-Trp and 7-Br-Trp serve as references. WT, wild type.

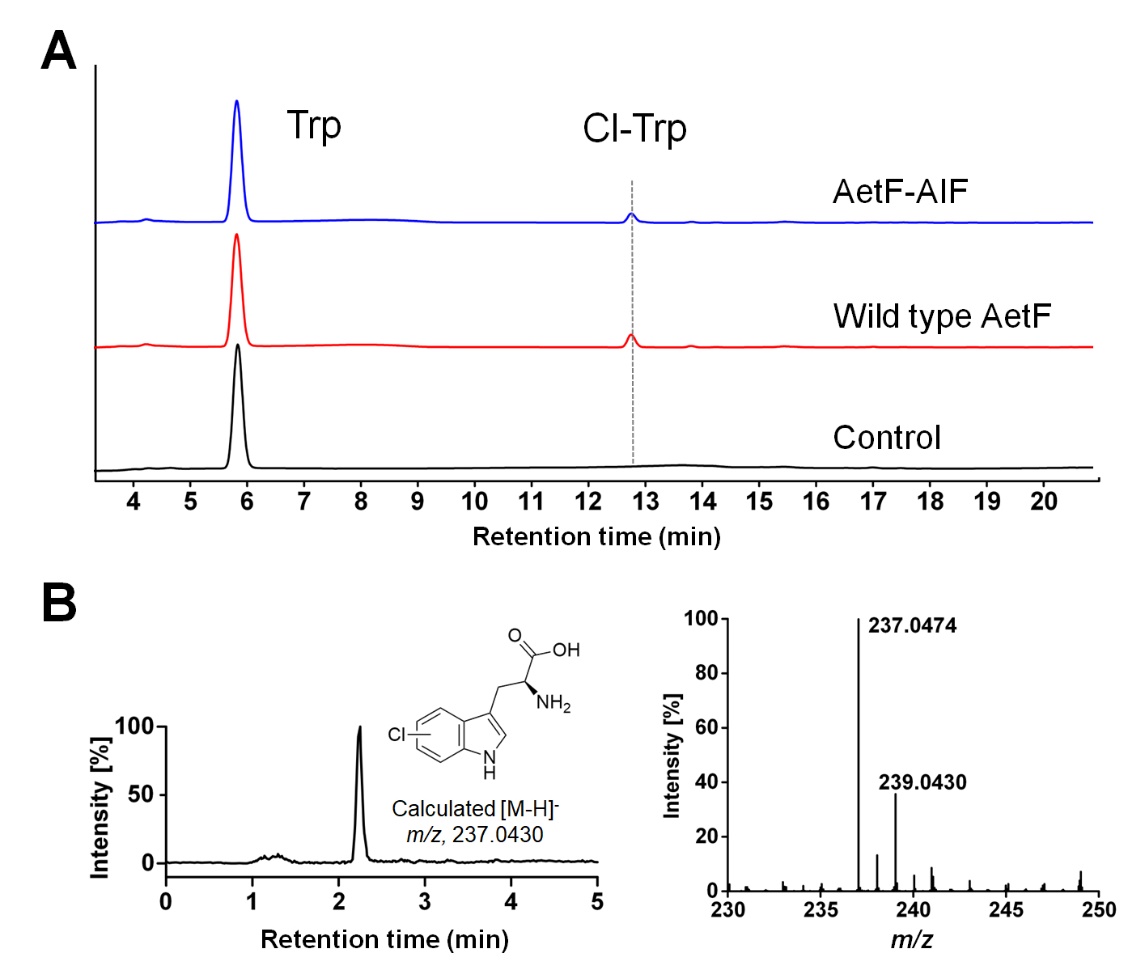

**Figure S5. The chlorination capacity of wild type AetF and AetF-AIF.** (**A**) The representative HPLC chromatograms (absorbance at 280 nm) of the product of wild type AetF- or AetF-AIF-catalyzed halogenation reactions that contain 500 mM sodium chloride and 6.5 μM enzyme. Reaction time, 5 hrs. (**B**) The extracted ion chromatogram and mass spectrum of chlorinated Trp detected in the AetF-catalyzed reaction shown in (**A**). The calculated monoisotropic [M-H]^-^ *m*/*z* value for the chlorinated Trp (^35^Cl) is displayed on the left panel.

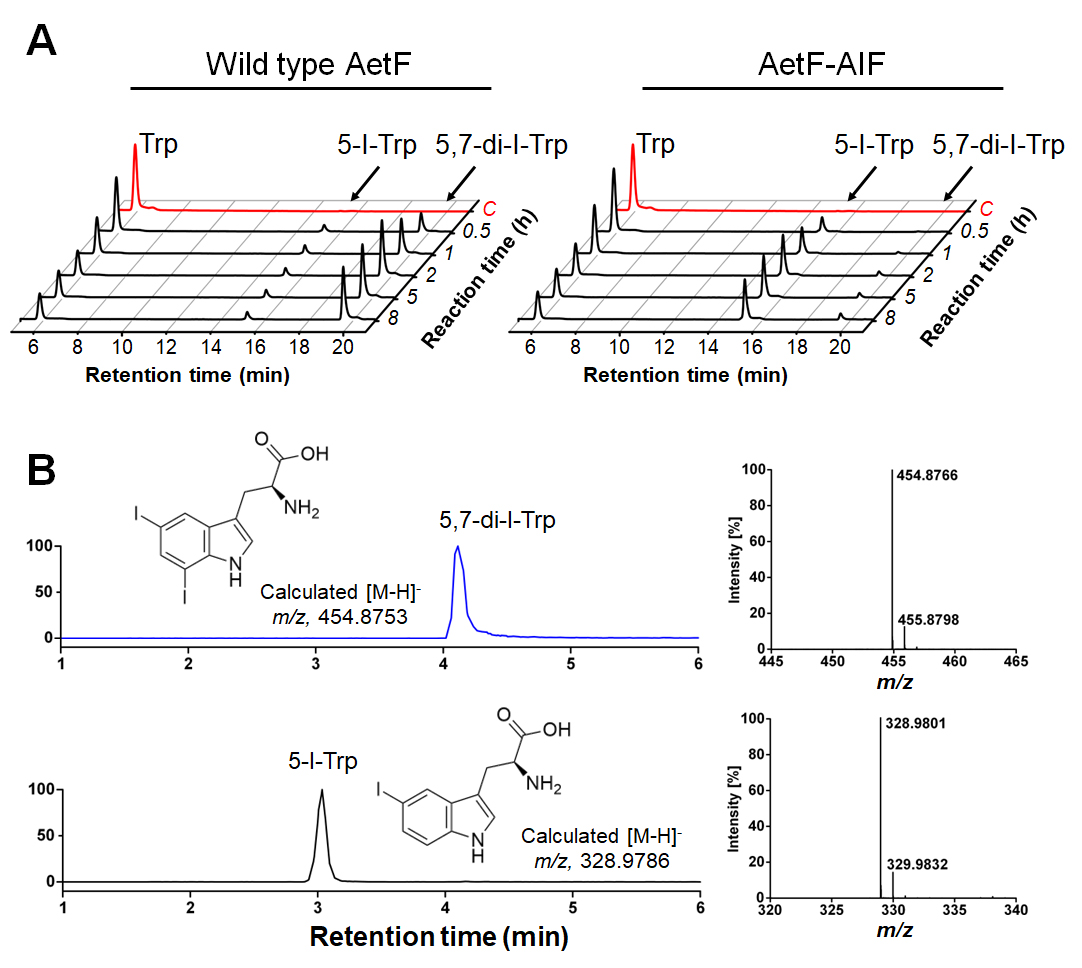

**Figure S6. The iodination capacity of wild type AetF and AetF-AIF.** (**A**) The representative HPLC chromatograms (absorbance at 280 nm) of products of wild type AetF- or AetF-AIF-catalyzed iodination reactions displayed in a reaction time-dependent manner. (**B**) The extracted ion chromatograms and mass spectra of 5-I-Trp and 5,7-di-I-Trp detected in the AetF-catalyzed reaction shown in (**A**). The calculated monoisotropic [M-H]^-^ *m*/*z* values for the displayed compounds are displayed on the left panel.

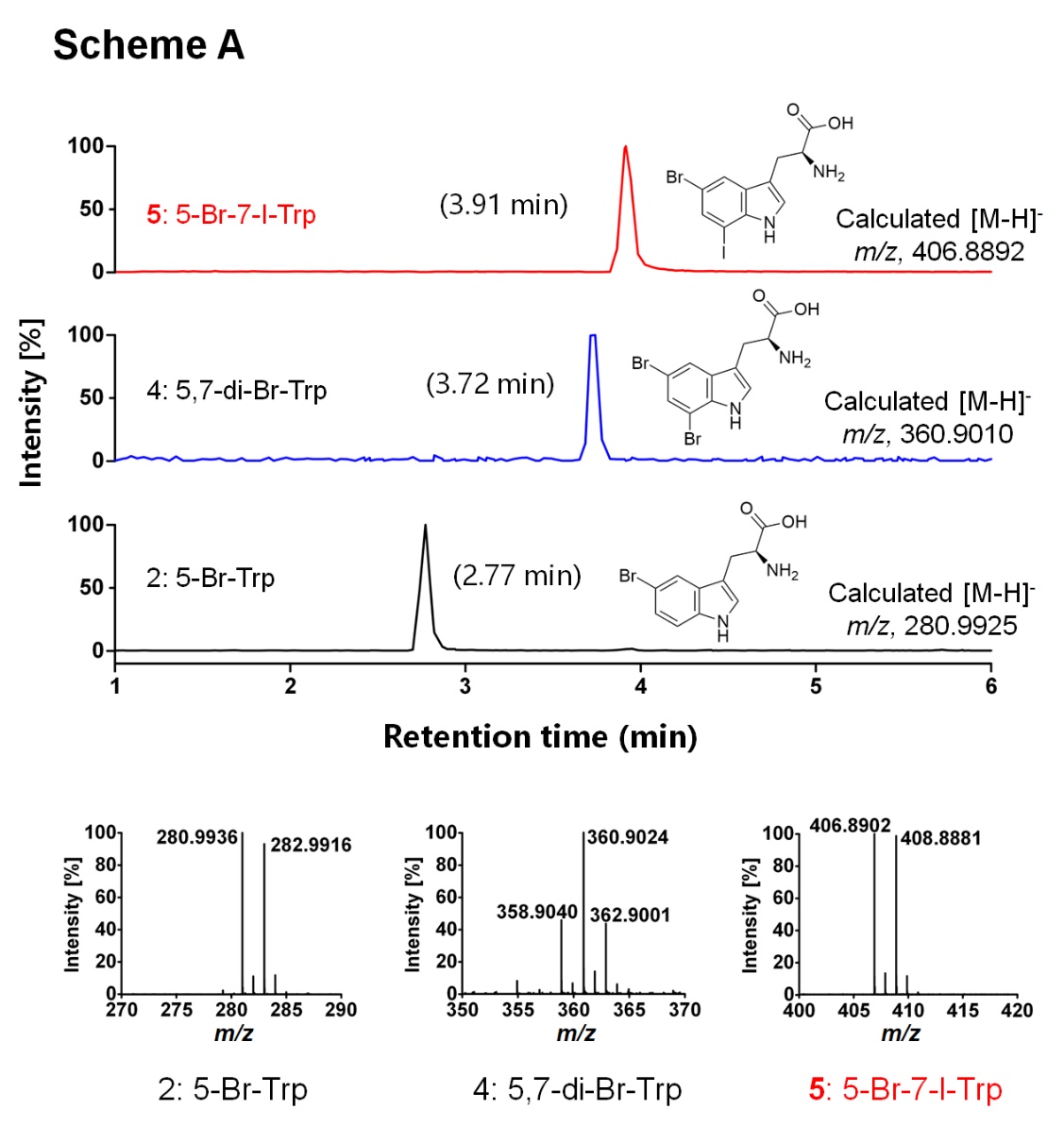

**Figure S7. Extracted ion chromatograms and mass spectra of each component in the reaction mixture from scheme A.** The calculated monoisotropic [M-H]^-^ *m*/*z* values for the displayed compounds (^79^Br for brominated compounds) are displayed aside.

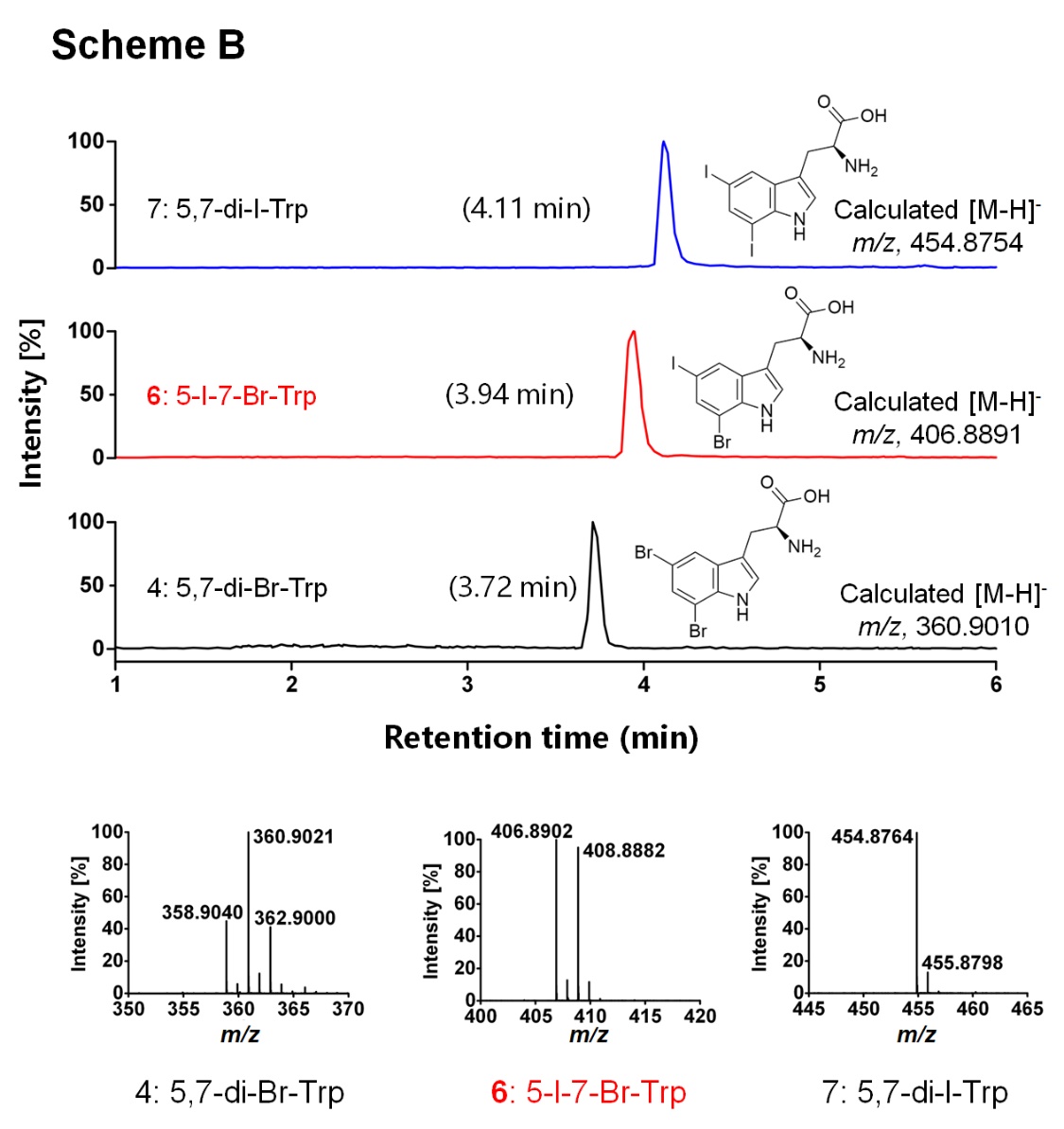

**Figure S8. Extracted ion chromatograms and mass spectra of each component in the reaction mixture from scheme B.** The calculated monoisotropic [M-H]^-^ *m*/*z* values for the displayed compounds (^79^Br for brominated compounds) are displayed aside.

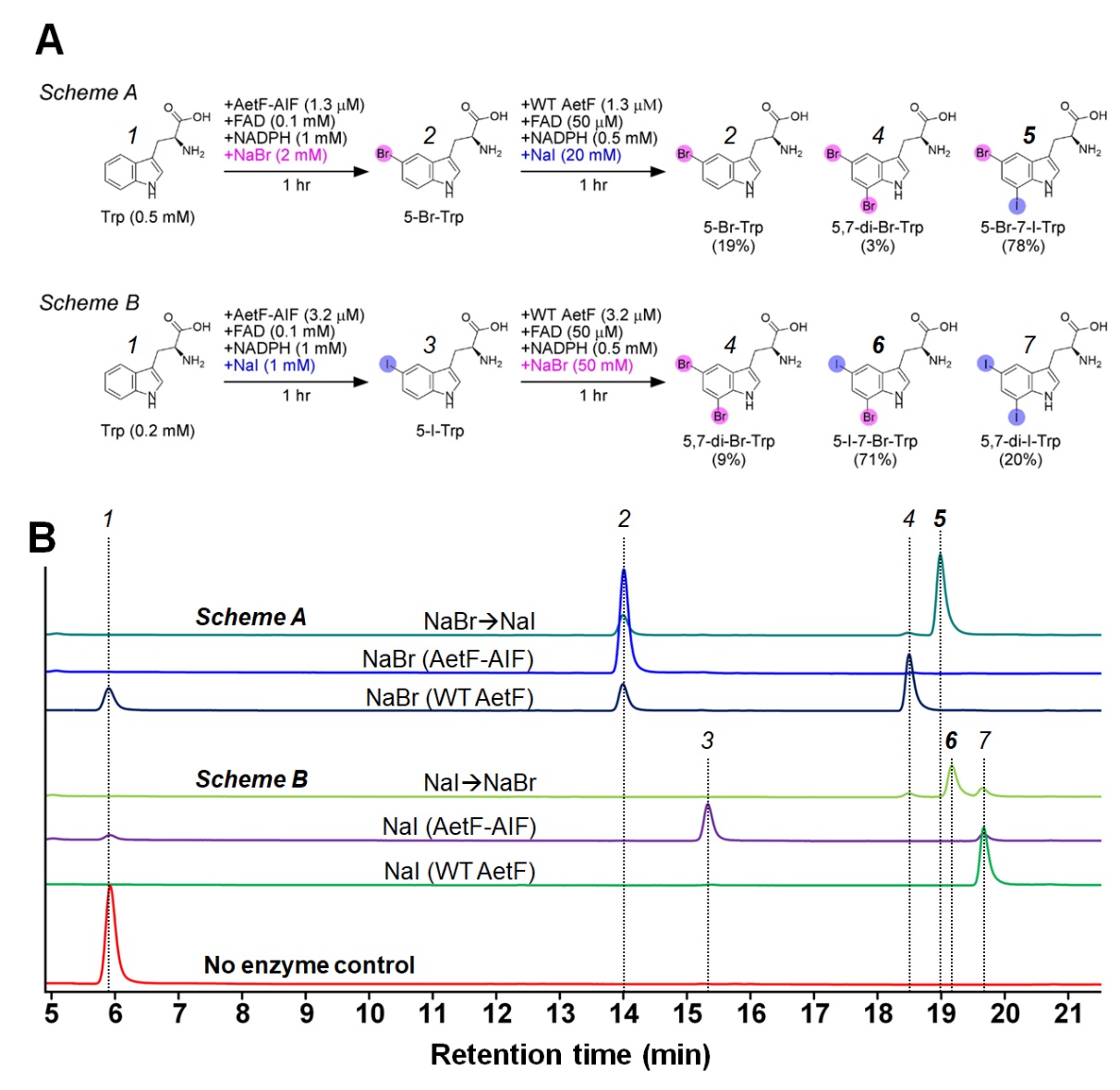

**Figure S9. Synthesis of heterogeneously di-halogenated Trp in 2-mL one-pot reactions.** (**A**) The designation and the composition of the final product of two schemes. (**B**) The representative HPLC chromatograms of reaction mixtures yielded from two schemes described in (**A**). 1, Trp; 2, 5-Br-Trp; 3, 5-I-Trp; 4, 5,7-di-Br-Trp; 5, 5-Br-7-I-Trp; 6, 5-I-7-Br-Trp; 7, 5,7-di-I-Trp.

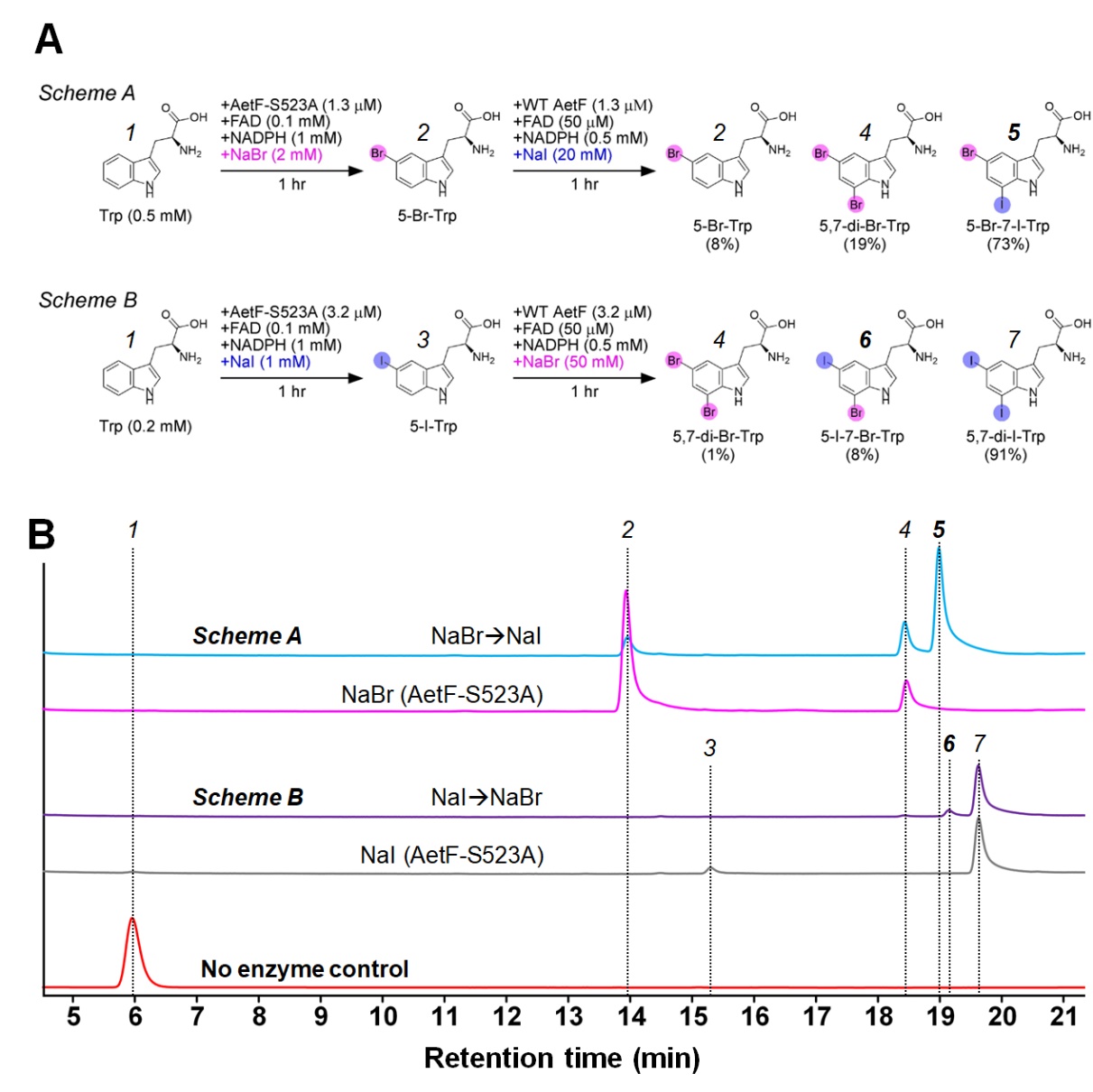

**Figure S10. Synthesis of heterogeneously di-halogenated Trp by using variant AetF-S523A.** (**A**) The designation and the composition of the final product of two schemes. (**B**) The representative HPLC chromatograms of reaction mixtures yielded from two schemes described in (**A**). 1, Trp; 2, 5-Br-Trp; 3, 5-I-Trp; 4, 5,7-di-Br-Trp; 5, 5-Br-7-I-Trp; 6, 5-I-7-Br-Trp; 7, 5,7-di-I-Trp.
